# Long-Read Low-Pass Sequencing for High-Resolution Trait Mapping

**DOI:** 10.1101/2025.01.09.632261

**Authors:** Kendall Lee, Walid Korani, Philip C. Bentz, Sameer Pokhrel, Peggy Ozias-Akins, Alex Harkess, Justin Vaughn, Josh Clevenger

## Abstract

Accelerating crop improvement is critical to meeting food security demands in a changing climate. Long-read sequencing offers advantages over short-reads in resolving structural variations (SVs) and aligning to complex genomes, but its high cost has limited adoption in breeding programs. Here we develop a high-throughput, scalable approach for long-read low-pass (LRLP) sequencing and variant analysis with PacBio HiFi reads, and apply it to trait mapping in a complex tetraploid peanut (*Arachis hypogaea*) genome multi-parent advanced generation intercross. We analyze LRLP using both a single reference genome and a pangraph, using both proprietary and open-source tools to analyze SVs and coverage. An increased number of variants are consistently called for LRLP data compared to short-read data. At 1.63x average depth, LRLP sequencing covered 55% of the genome and 58% of gene space, outperforming 1.68x depth short-read low-pass sequencing, which achieved only 17% and 11%, respectively. Enhanced data retention after filtering for probabilistic misalignment and an ∼8.5x decrease in cost per value further demonstrated LRLP’s efficacy. Our results highlight LRLP sequencing as a scalable, cost-effective tool for high-resolution trait mapping, with transformative potential for plant breeding and broader genomic applications.

## Introduction

The global population is projected to approach 10 billion by 2050 (UN Population Division, 2024). Meeting the caloric needs of this growing population will require a 60% increase in agricultural crop production (1). Compounding this challenge, climate change has already reduced global crop yields by 21% since the 1960s, presenting additional hurdles for farmers and plant breeders (2). Addressing these challenges necessitates rapid crop improvement to enhance yield and productivity while overcoming climate-induced stresses such as drought and disease.

One significant obstacle in plant breeding is the time required to evaluate new cultivars for desirable traits, particularly in perennial crops where traits may take years to phenotype. However, advances in DNA sequencing technologies over the past decade have paved the way for molecular-based plant breeding. Marker-trait association techniques like Genome-Wide Association Studies (GWAS) and Quantitative Trait Locus sequencing (QTL-seq) have accelerated the development of new cultivars by enabling trait-marker-based selection (3–5). These approaches primarily rely on short-read sequencing for trait mapping, which involves statistical analyses to correlate genetic variants with phenotypic traits (6, 7). Short-read sequencing remains popular due to its affordability, with low-pass sequencing further driving down costs. Studies have demonstrated the effectiveness of short-read low-pass sequencing for genomic selection in animals (8, 9). Khufu, a proprietary single reference genotyping analyses, has achieved 100% SNP calling accuracy with depths as low as 0.5x in plants (10, 11). However, the short-reads have significant limitations (12, 13). Typically generating read lengths of around 150 base pairs (bp), short-reads often align ambiguously to multiple regions of a reference genome, reducing confidence in variant calls and filtering out potentially informative data. This is especially problematic in polyploids. Moreover, short-read sequencing fails to capture large structural variations (SVs) such as insertions, deletions, inversions, translocations, and duplications, which are increasingly recognized as key contributors to agronomic traits (14). For instance, SVs have been implicated in important traits in melon, lettuce, peach, and rapeseed (15–19).

Long-read sequencing has emerged as a powerful tool for reference genome assembly, capable of generating sequences tens of thousands of basepairs in length (20). These longer reads simplify alignment to reference genomes and enable the detection of SVs (21). Despite these advantages, long-read sequencing has historically been limited to high-coverage sequencing of a small number of samples due to the high cost and low throughput of wet lab protocols for extracting high molecular weight DNA and preparing sequencing libraries (22). However, our development of long-read low-pass (LRLP) sequencing offers a promising solution. By pooling multiple samples for sequencing at low coverage, LRLP significantly reduces the cost per sample while preserving the advantages of long reads for variant calling. This makes long-read sequencing feasible for routine integration into breeding programs.

Pangenomic graphs, which incorporate multiple reference genomes to capture broader species diversity, improve variant detection in complex genomes (23–25). These graphs consist of a core set of homologous regions across genomes, as well as divergent regions, offering a more comprehensive framework for trait mapping.

Peanuts (Arachis hypogaea) are a challenging allotetraploid crop for trait mapping due to their genomic complexity (26). Multi-parent advanced generation intercross (MAGIC) populations provide an effective framework for trait mapping, offering enhanced resolution and genetic diversity (27). These populations are particularly valuable for incorporating multiple traits of interest into breeding programs (28).

We present the efficacy of the LRLP methodology within a functional peanut breeding framework. Using an 18-parent (16-way) MAGIC peanut population, we demonstrate the development of wet lab tools and the adaptation of the Khufu analysis pipeline (10) to identify and associate SNPs and SVs with traits of interest. By aligning both LRLP and short-read sequences to a single reference graph and the pangraph of the 18 parents, we evaluate the performance and utility of each approach. We demonstrate that LRLP offers a robust and highly accurate alternative to current methods of selecting lines with QTLs of interest in an active peanut breeding program.

## Results

### Sequencing

We sequenced individuals from an active peanut breeding population to show the applicability of LRLP, a group of individuals that came from 18-cross hybrids then backcrossed with the intent to make selections using the variant data found. The population segregates for six known QTL that can be selected for using genomic information.

Sequencing statistics are reported from PacBio’s SMRTLink software. These statistics are important to understand the efficacy of our sample preparation pipeline. Pools consisted of 10 to 13 samples per pool with the majority having 12. Overall yield per sample, base quality score, mean read length, and median read length were evaluated to ensure high-quality sequences were being produced. Out of 136 samples, nine were removed due to low sequencing yield. An average of 4.25 GB of HiFi sequence data was produced per sample and an average of 98.2% of HiFi basepairs across all samples had a Phred score above 20, showing that the quality of the data produced was very high (Table 1). Mean read length varied between 4,151 bp and 13,169 bp with an average of 7,927 bp (Table 1). Median read length had a similar range of 2,278 bp to 13,185 bp with an average of 7,046 bp (Table 1). Overall, high-quality data was produced using a high-throughput DNA extraction and library preparation protocol-a massive achievement for plant DNA.

**Table 1.**
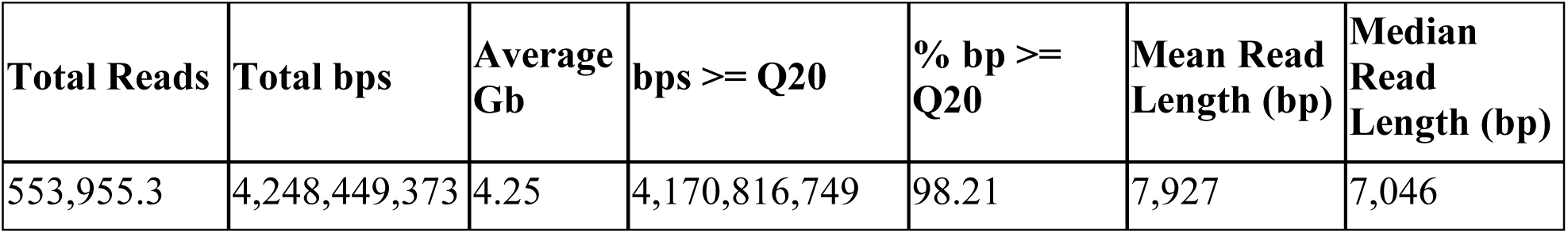
Raw sequencing statistics of long-read low-pass data of 127 MAGIC-population peanut lines.

### Coverage

Depth or depth of coverage is the average number of times a position in the genome has been sequenced, whereas coverage refers to the percentage of the entire genome that has been sequenced at least once. Both metrics were calculated for each sample. While depth was not normalized on a per-sample basis, a paired t-test shows that there is no significant difference between the long and short-read sequence depths (p-value = 0.8237), and the average difference in depth across samples is 0.028x (Fig. 1C). The average depth for long reads was 1.63x and the average coverage was 71.29%, whereas for short-reads it was 1.68x depth and 32.67% (Fig. 1D). After filtering for highly probable misalignments, long-read samples had an average coverage of 55% and short-reads had an average coverage of 17.3%. While the difference in depth was not statistically different for short and long reads, the difference in coverage and filtered coverage were both significantly different (p < 2.2×10^−16^ for both) (Fig. 1). The gene space (the portion of the genome containing genes) covered was also analyzed using the gene annotation for the reference genome, Tifrunner version 2. The average gene space covered for long reads after filtering was 57.9%, slightly higher than the overall filtered coverage (Fig. 1A & 1B). This is likely due to genic regions having greater sequence complexity. In contrast, after filtering, short-reads only cover 11% of gene space on average and are significantly lower than long reads (p < 2.2×10^−16^). Short-reads are ∼150 bp long and therefore are not long enough to cover genes appropriately. Also, high throughput short-read libraries are often prepared with PCR which may result in amplification of only some regions leading to unevenly distributed sequencing coverage. This significant increase in genome and gene coverage that LRLP sequences have even at the same depth as short-reads creates an immense advantage for variant calling for trait discovery. Far more of the genome is represented allowing for an increase in variants called. Moreover, an increase in gene coverage allows for more impactful variants to be discovered.

**Figure 1.**
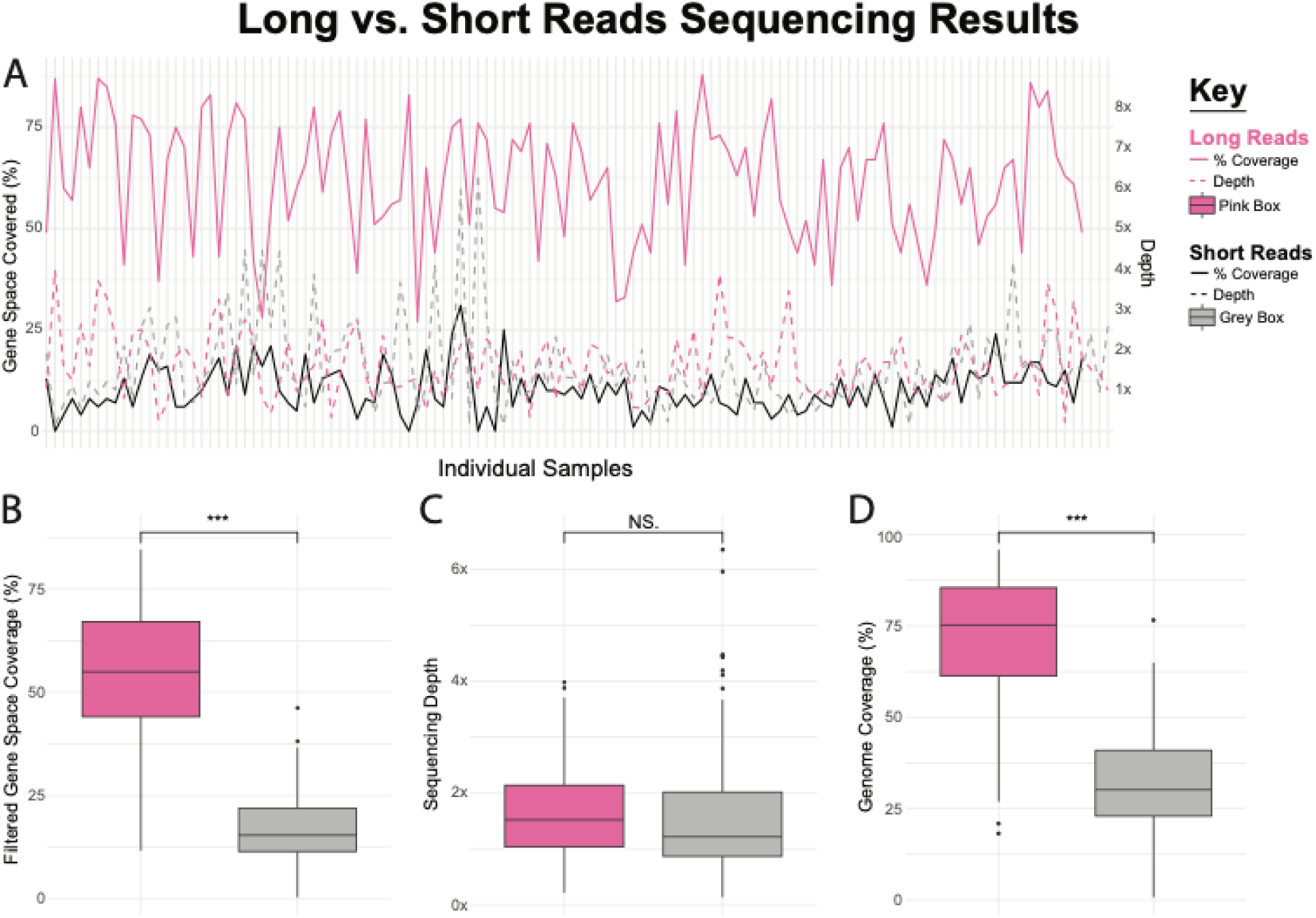
A) Line plot comparing sequencing performance for individual samples. Solid lines represent percent of genespace covered and dashed lines represent sequencing depth. Short-read sequencing is shown in black, while long-reaad sequencing is shown in pink. B) Box plot comparing filtered gene space coverage (%) between long-read (pink) and short-read (gray) sequencing. The long-read approach shows significantly higher coverage. (C) Box plot comparing sequencing depth between long-read (pink) and short-read (gray) sequencing. No significant difference (NS) is observed. (D) Box plot comparing genome coverage (%) between long-read (pink) and short-read (gray) sequencing. Long-read sequencing achieves significantly higher genome coverage. *** indicates statistical significance (p < 0.001). NS indicates no statistical significance.

### Confidence of Accurate Alignment

One of the key advantages of long-read sequencing over short-read sequencing is its higher likelihood of uniquely aligning to the reference genome, resulting in fewer sequences being discarded due to multiple alignments. We refer to the percentage of reads retained after filtering for probability of misalignment as confidence of alignment. Misalignment filtering is done when a read maps to a different region of the genome more than once. On average, long reads had a confidence of alignment of 92.9%, significantly higher than the 54.9% retention rate observed for short-reads (p-value < 2.2e-16) (Supp. Fig. 1). This increased alignment confidence in long reads enables more comprehensive genome coverage, especially in repetitive or complex regions that are challenging for short-reads to resolve.

### Total Calls

Against a single reference genome, long reads with Khufu called a total of 235,583 SNPs after applying several filters: quality control, uniquely mapping reads, at least 1x depth, minor-allele frequency threshold (SNPs with a value lower than 10% are dropped), and removing missing data across SNPs (SNPs with calls missing from 75% or more of the lines). Linear Khufu with short-reads called only 35% of the same number of calls under the same filtering parameters. The drastic increase in SNPs found increases the likelihood of correctly correlating a SNP to a trait of interest.

### Imputation Accuracy

Khufu fills in missing SNPs using a reference-panel-free imputation method. The khufu pipeline has a built-in check to ensure that imputation calls are correct. It masks some percentage of the genome and makes imputations with the remaining data. It then checks the accuracy of imputations after unmasking the data. We find that despite almost 3x the number of SNPs called in LRLP data, the imputation accuracy remained similar. Linear khufu reports imputation accuracy at 25% genome-masked, 50% genome-masked, and 75% genome-masked. The long read low pass data increased from 89.2% accurate at 75% masked to 96.2% accurate at 25% masked. Short-read linear khufu data showed similar imputation accuracy ranging from 90% at 75% masked to 96.69% at 25% masked.

### Structural Variation and Impact

Large SVs are mostly missed by short-read mapping as most short-reads are 150 bp after sequencing and more like 135 bp after adapter trimming. LRLP does not have the same problem since the read length is often long enough to span SVs. We called SVs in the samples using the pbsv pipeline (29) with TRv2 as the reference genome. The total number of unique SVs reported across all lines was 107,178, with insertions and deletions being the most numerous. The average insertion length was 173.74 bp and the average deletion length was 1246.35 bp across all samples, both higher than the short-read length of ∼150 bp. The software program snpEff was used to evaluate the potential effects of the SVs (Cingolani et al., 2012). SnpEff reports types of SV variation as well as the potential impact. The impact levels are high, moderate, low, and modifier. The ‘high’ impact SVs are predicted to be very disruptive to protein coding and are mutations such as frameshift and nonsense mutations. ‘Moderate’ impact SVs would not disrupt the protein but may change its effectiveness. These are mutations such as missense variation and inframe deletions. ‘Low’ impact variants are unlikely to result in any change. These are mutations such as a synonymous variant. ‘Modifier’ variants are a class of impact non-coding variants or variants affecting non-coding genes. These are cases where predictions are challenging and/or there is no evidence of impact. Sample vcf files were merged to evaluate the unique variant’s impact. 1,221 high impact, 464 moderate impacts, 252 low impact, and 96,767 modifier impact unique variants were found across all lines. Of the high-impact variants, 304 were insertions, 843 were deletions, 2 were breakpoints, 110 were duplications, and 14 were inversions.

### Utilization of pangenome graphs to maximize genotyping

To maximize the efficiency of profiling genotype information with long read sequencing, we developed a pipeline (KhufuPAN) that utilizes vg tools/giraffe to align the long reads and call high quality variants (30). After processing the graph and maintaining high quality variants, long reads are aligned to the processed graph using giraffe. Variants are aligned to the high quality variant set, and missing data and minor allele frequency is calculated for customized filtering. Finally, a custom file format is generated we call a panmap. The panmap includes critical information for each variant type, including chromosome and position, length of alleles, alleles assigned to each member of the graph, and then genotype calls in numerical format. Along with the panmap file is a fasta file with each allele. The panmap format can be converted to hapmap or vcf for use in different programs, making the output of KhufuPAN informative and flexible. Further, the speed of analysis makes KhufuPAN very efficient. The current experiment used an average of 47.9 GB maximum RAM and 24.15 CPU hours for completion per line with the majority of the pipeline supporting multiple threads, increasing efficiency.

### PanGraph Alignment

A 16-genome pangraph was constructed with the parental genomes of the sample lines and the long read low pass sequences of the sample lines were aligned to it. The pangraph captured 3,042,561 variants for long reads while it captured 186,509 for short-reads, only 6% of the number that long reads captured. While there was no significant difference between the percentage of missing data for short-reads vs long reads on the pangraph, LRLP sequences identified significantly more variation (Fig. 2, Supp. Fig. 4). On average, short-reads identified 76,866 SNPs, 13,228 indels and 15 large SVs per line, while long reads found 1,436,345 SNPs, 606,720 indels (2 to 1000 bp), and 774 large SVs (>1000 bp) per line (Fig. 2). Long reads identified approximately 18.7x more SNPs, 45.9x more indels, and 51.6x more SVs compared to short-reads. To further parse the SVs found we used a log scale to group the variants by length spanning from 1 bp to ∼50kb and find that the LRLP data is significantly superior in capturing more and longer SVs in every length group (Supp. Fig. 5). This highlights the wealth of information that LRLP sequences offer in comparison to short-reads.

**Figure 2.**
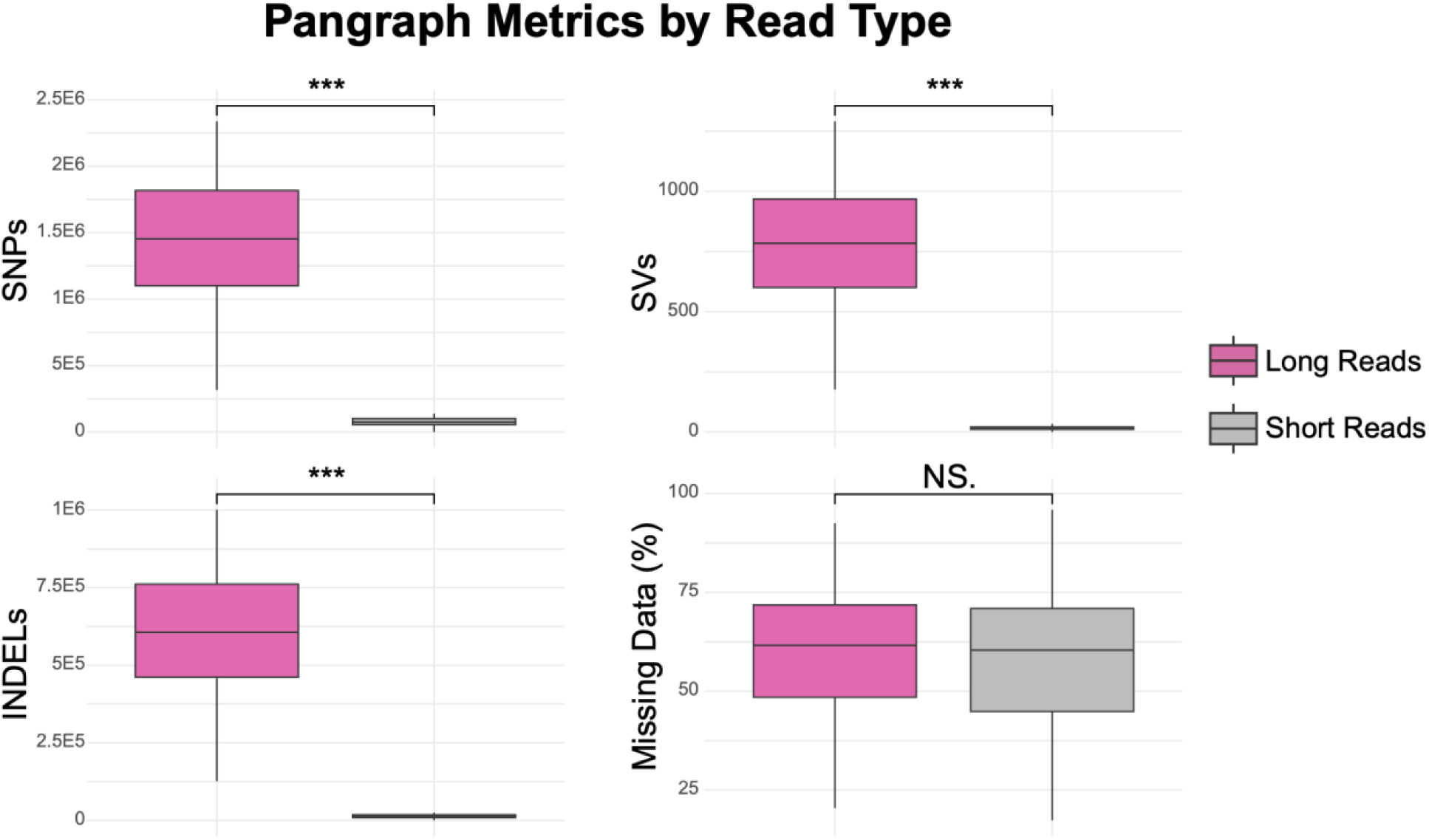
Boxplots comparing the performance of low-pass long and short-reads across different variants as called on the dynamic pangraph. Long read analysis finds significantly more SNPs, indels and large SVs than short-reads. There is no significant difference between the rate of missing data between long read and short-reads. The asterisks (***) indicate significant differences between long and short-reads for the respective metric. Indels are defined as variants between 2 and 1,000 bp in length. Large SVs are defined as variants above 1,000 bp in length.

### Selection Potential

Accurately finding and selecting lines that actually have the QTL of interest is the main challenge for plant breeders in this era of genomic selection. Typically SNP markers are used for this purpose. However low density markers can miss causative variants or regions closely linked to the QTL of interest, reducing the accuracy of selection. This is particularly problematic in regions of the genome with high recombination rates or complex structural variation, where linkage between markers and causative genes can break down. Additionally, reliance on SNPs that are not functionally validated may lead to the selection of markers that are merely correlated with the trait in a specific population, rather than directly influencing the phenotype. This can result in false positives or missed opportunities to select for beneficial alleles, ultimately limiting the effectiveness of breeding programs. Whole-genome sequencing (WGS) with short-reads is better for enabling the accurate detection of SNPs and small indels across the genome, which facilitates genomic selection. However, short-reads struggle to resolve complex genomic regions and capture SVs. LRLP sequencing offers the best solution. While the lower coverage reduces sequencing costs, the long reads enable superior genome mapping, capturing structural variants and resolving complex regions. This comprehensive view of the genome is particularly valuable for resolving complex QTLs and improving selection accuracy in plant breeding programs.

To test whether the immense amount of sequence information found with LRLP is useful in an applied way for plant breeding, we used a breeding population of peanuts to generate similarity scores for QTL regions of interest between the breeding population lines and the parents that are known to have the QTL region of interest. We find that regardless of depth, the LRLP sequences consistently have significantly higher locus similarity scores for important disease resistance loci for late leaf spot (LLS) and tomato spotted wilt virus (TSWV) (Fig. 3). This is likely due to the missing data in the region for short-reads. Further investigation into the TSWV locus shows the 19kb insertion is present as a single long read but not represented by the short-reads, further highlighting the efficacy of LRLP at capturing significant variation (Fig. 4).

**Figure 3.**
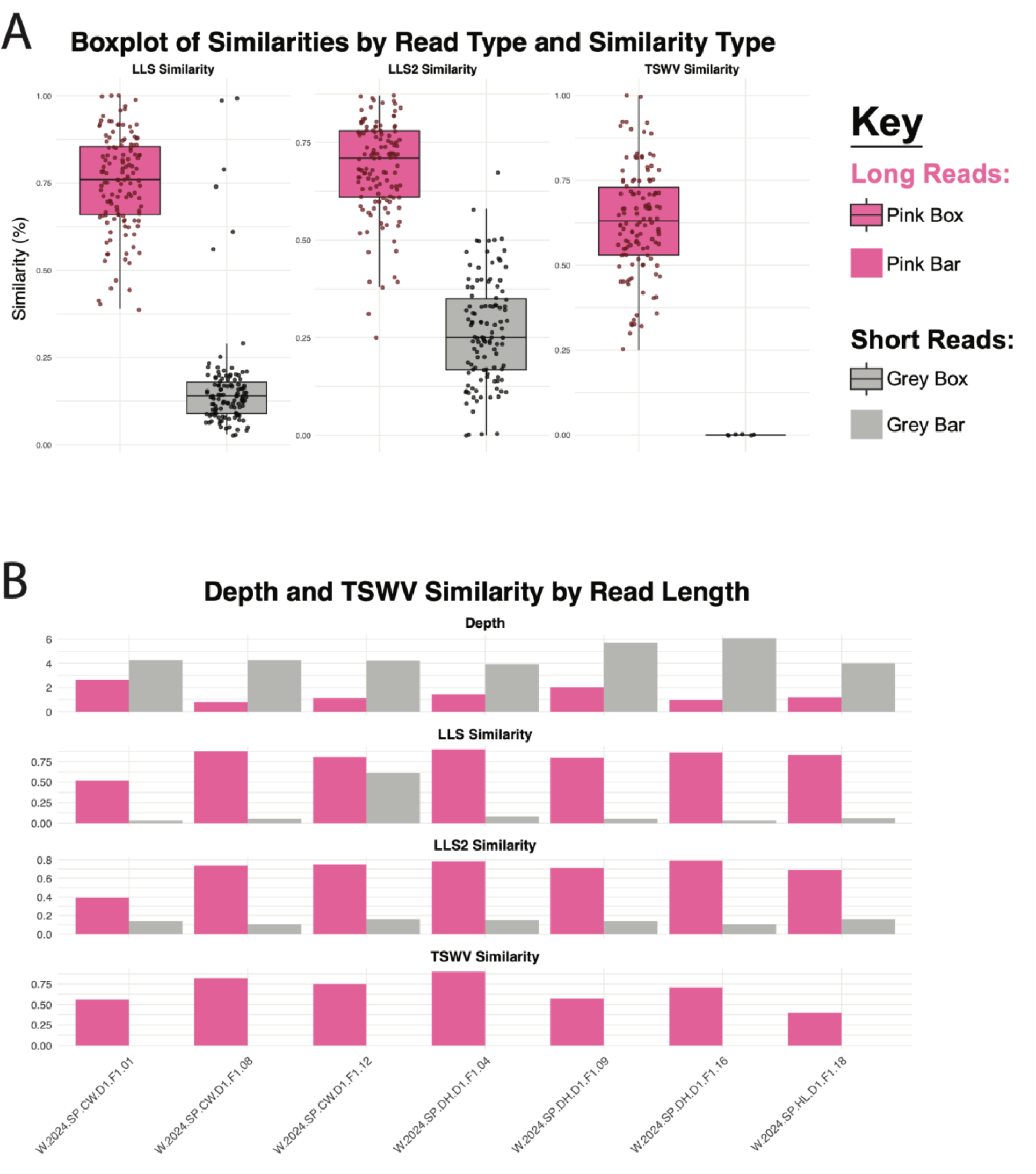
A) Boxplots showing similarity scores between long-read and short-read sequencing for loci for late leaf spot (LLS & LLS2) and tomato spotted wilt virus (TSWV). Each boxplot represents the distribution of similarity scores, with long reads shown in pink and short-reads in gray. Points represent individual sample measurements, and boxplots highlight the median and interquartile range. B) Bar plots illustrating depth and trait loci similarity scores by sample ID for long-read and short-read sequencing, where the short-reads have a much higher depth than the corresponding long

**Figure 4.**
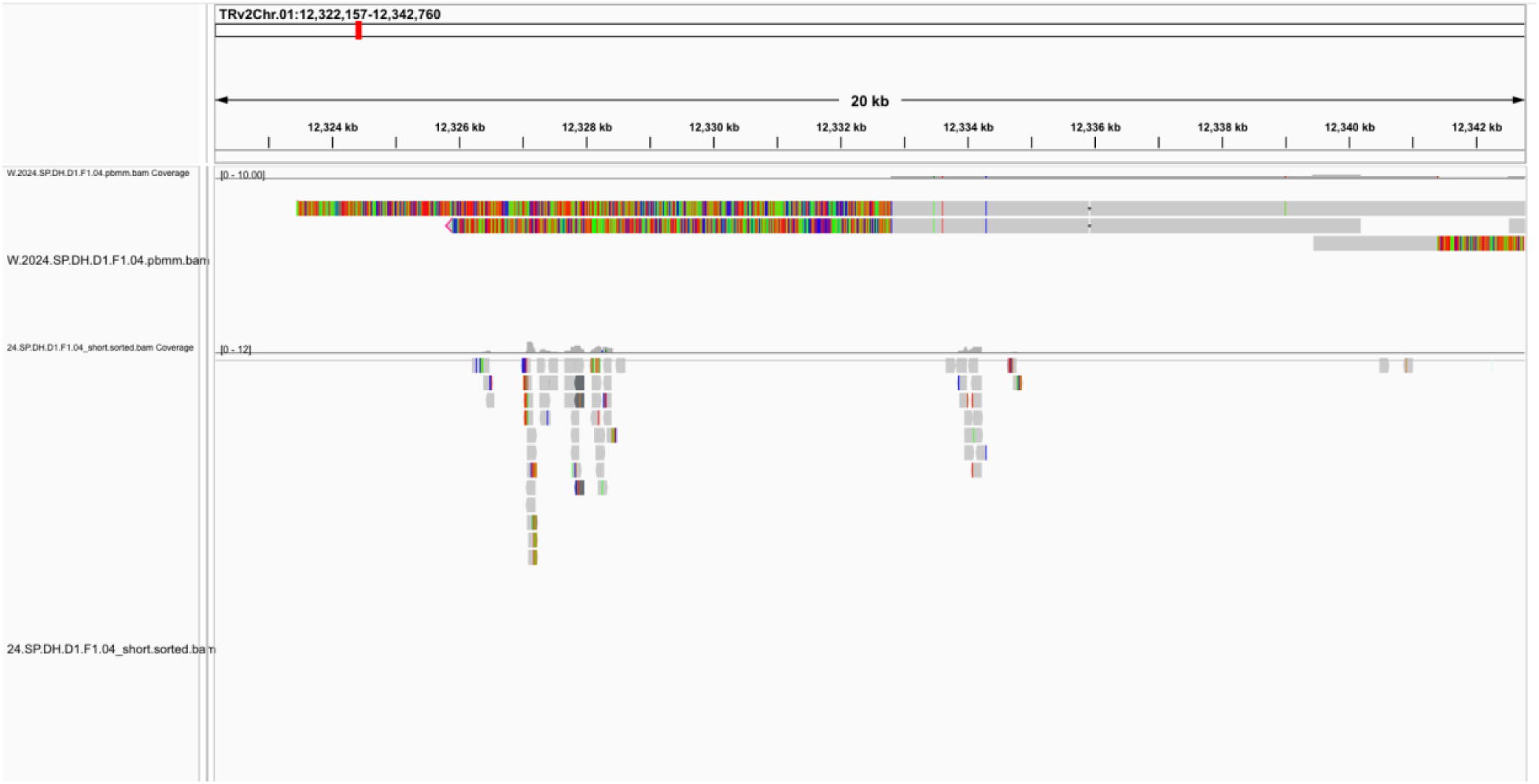
An integrated genomics viewer (IGV) screenshot alignment of long and short-reads to the region of TRv2 where the large insertion that confers resistance to TSWV maps. TRv2 contains one glutamate receptor gene in the region but lack the multiple copies that are found in the insertion, which confer resistance. The LRLP reads are on the top and show a long read with clipped reads spanning a 20kb region, covering the entire region. The partial clipping of the read indicates the insertion. Short-reads are on the bottom and show some mapping to the region, but not enough to discover the TSWV disease resistance region insertion. Short-ads are mapping to the edge of the insertion to one gene that is found in TRv2 but does not pick up on the additional genes which make up the insertion.

### Cost Per Insight

A comparison of sequencing approaches highlights the cost-effectiveness of long-read sequencing in achieving comprehensive genomic coverage. Despite the higher initial investment, long reads consistently deliver superior coverage across filtered gene space and the entire genome compared to short-reads, as shown in Figure 1. The cost per insight analysis (Fig. 5) reinforces this efficiency, demonstrating that long reads provide significantly greater genomic insight per unit cost. Calculated as the cost per insight for short-reads divided by that for long reads, the fold change highlights the superior cost-effectiveness of long reads. Across all samples, the average fold change is 8.53, indicating that long reads deliver over eight times the cost efficiency of short-reads. This substantial difference reinforces the economic advantage of long-read sequencing for projects requiring high-resolution genomic data. While this cost difference is due to the larger number of variants the LRLP data provide, we show in the previous section (Fig. 3 and 4) that these extra variants are necessary for accurate selection. These findings position long-read sequencing as a more economical choice for high-resolution genomic analyses, particularly in projects requiring extensive coverage of complex genomes.

**Figure 5.**
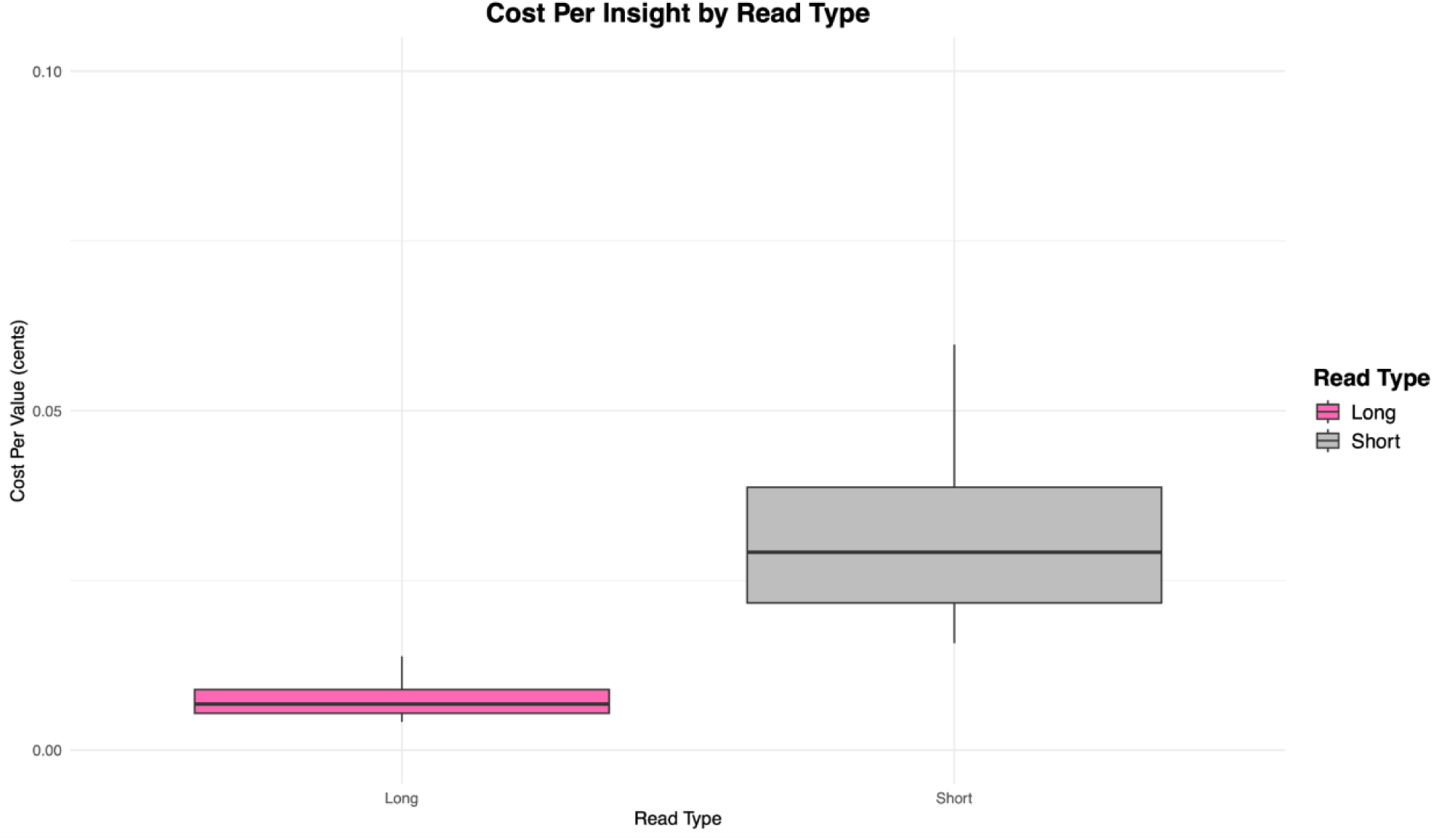
A boxplot of cost per value for LRLP sequencing versus low coverage short-read sequencing when genotyping using a pangenome graph as a reference. Using a cost calculation based on current prices for the kits, consumables, and services for the Clevenger Lab in Huntsville, Alabama in the year 2024, LRLP cost 8.53 times less than the short-read counterpart from an insight (variant) standpoint.

## Discussion

Long-read sequencing is widely recognized across various fields for its superior ability to capture complex genetic variation compared to short-read sequencing (31), (32), (33), (21) and (34). Despite the increasing adoption of low-depth short-read sequencing, there has been hesitancy to explore LRLP sequencing, primarily due to challenges in high-throughput sample preparation. Extracting high molecular weight DNA remains a bottleneck, as most protocols are low throughput, processing only one or two samples at a time. However, establishing efficient methodologies for long-read genotyping is essential for advancing plant and animal breeding, as well as human clinical applications.

Currently, short-read sequencing dominates population genotyping due to its scalability in the wet lab and relatively low cost. However, the limited genomic resolution of short-reads often leads to missed key data, particularly SVs. Additionally, the short-read length increases misalignment rates, resulting in significant data loss during filtering. While short-reads remain valuable for single nucleotide polymorphism (SNP) genotyping, population-based long-read sequencing is gaining traction as a means to capture the broader genomic landscape.

Previous studies on low-depth long-read sequencing have been limited in scope, examining small sample sets. For instance, Malmberg et al. (2019)(35) used Oxford Nanopore Technology’s MinION platform to sequence nine Brassica napus samples at 2x to 5x depth, achieving accurate genotyping and improved alignment despite a 10% error rate. Our study builds on this by scaling up the sample size and leveraging PacBio HiFi sequencing, which offers significantly lower error rates (reads > Q20).

We developed a high-throughput methodology for DNA extraction and library preparation using new PacBio reagents and kits on the Revio sequencer. Our optimized protocol produces medium-quality DNA at lower concentrations compared to traditional low-throughput methods. However, subsequent steps—including short-read elimination, bead-based cleanups, and sample pooling—address these limitations. Targeting a depth of 2–3x, our samples achieved an average depth of 1.63x, with variation between 0.5x and 4x, largely attributed to differences in sequencing yield (r = 0.93, p < 2.2×10⁻^16^).

This study demonstrates that low-depth long-read sequencing can effectively capture genomic data at scale. Our results align with findings from Malmberg et al., (2019) (35), with improved accuracy due to reduced error rates. Furthermore, the absence of PCR amplification in long-read workflows eliminates bias, a key advantage over short-read methods. Magner et al. (2024) (36) recently combined 4x long-read data with short-read sequencing and found enhanced SNP detection and large SV calling, further highlighting the value of integrating long reads into genotyping pipelines. While more studies are necessary to bolster findings, the results of our work and the few others who have explored variations of LRLP clearly show the utility of this method.

This research and method development shows that long-read HiFi sequencing can be scaled up for direct use in breeding programs for routine selections. The cost of LRLP largely depends on the genome size of the organism of interest, but regardless, it is much more affordable than traditional high-coverage long-read sequencing. LRLP could be used in a variety of ways to improve a breeding program. Breeders who may choose not to fully convert to LRLP for every generation could strategically implement it in key pieces of the breeding cycle. For example, a breeder may choose to use short-read skim-seq early on and then LRLP for late-generation testing because there are less lines to test and the accuracy for selection is much better (Fig. 3). On the other hand, they could also use LRLP to capture all the genetic information on their founder lines before initial crossing as a cost effective alternative to genome assembly. LRLP could also be used just once as an improved alternative to short-read whole genome sequencing for SNP panel development. Overall, LRLP offers a versatile and highly accurate tool to improve genomic selection in plants and animals.

The potential applications of LRLP extend beyond plant and animal breeding. In microbial genomics, where genomes are smaller, LRLP offers cost-effective reference-based sequencing with enhanced resolution, improved taxonomic analysis, and more accurate variant calling (37). This versatility positions LRLP as a transformative tool across multiple fields.

### Conclusion

Our study demonstrates that long-read low-pass sequencing (LRLP) is a powerful, cost-effective alternative to short-read sequencing for high-resolution trait mapping in peanut breeding. By leveraging LRLP, we achieved significant improvements in variant detection, particularly for structural variations that are often missed by short-read methods, and show the necessity of these extra variants in accurately selecting for QTL of interest to our peanut breeding program. The combination of LRLP with pangenomic approaches provides a scalable solution for accelerating genomic analyses in breeding programs. As sequencing costs continue to decrease, LRLP sequencing holds extraordinary potential for transforming plant and animal breeding efforts, enabling more precise and efficient trait selection across diverse species.

## Online Methods

### Population Development

The peanut population tested in this study are from an active breeding program from HudsonAlpha Institute for Biotechnology and the University of Georgia. The goal of the breeding program is to stack known desirable QTL from parents into progeny. Eighteen divergent parent lines were crossed in pairs and seven F1 plants per cross were kept from the original crossing in January of 2019. A half-diallel cross was then done in September of 2020 with the progeny of the first cross. Seventy-five 4-way crosses were carried out with the progeny in March of 2021 and finally, fifty-six 8-way crosses were carried out to combine the genotypes of 18 parents in September of 2021. These crosses were then selfed to produce F2 seeds.

### DNA Extraction and Library Preparation

Approximately 100 mg of young leaf tissue was collected from each individual in the greenhouse using a hole punch and immediately transferred into pre-cooled 2 mL deep-well 96-tube Eppendorf plates containing grinding beads. The plates were kept on dry ice during collection and stored overnight at -80°C. Tissue grinding was performed using a Spex MiniPrep G in short intervals, allowing the samples to re-freeze on dry ice between cycles to maintain tissue integrity.

For long-read sequencing, DNA was extracted using PacBio-formulated reagents. Extractions were performed using either a full 96-well plate or subsets of 24 tubes. Whole-plate extractions were automated on a Thermo Scientific™ KingFisher™ Apex magnetic particle processor (cat. #5400930). DNA concentrations were measured using a Qubit™ High Sensitivity Assay Kit (cat. #Q33230). DNA fragments smaller than 10 kb were removed using the Pacific Biosciences Short-Read Elimination (SRE) kit (cat. #102-208-300), resulting in a typical recovery of approximately 50% of the original DNA concentration.

Following SRE, DNA was diluted to ≤10 ng/µL in 300 µL of low TE buffer. Shearing was performed using a Hamilton MicroLab Prep via repeated pipetting cycles. The DNA was recaptured in reduced volumes and cleaned with PacBio SMRTbell® Cleanup Beads (1:1 volume-to-volume ratio). Library preparation was carried out with the PacBio HiFi SMRTbell® Prep Kit 96 (cat. #103-381-200). Libraries were pooled with distinct adapter sequences for multiplexing, with pools typically containing 12 samples (ranging from 9 to 14 samples per pool). Sequencing was performed on PacBio Revio™ Flow Cells at SeqCenter (Pittsburgh, PA) using a PacBio Revio™ Sequencer.

For short-read sequencing, DNA was extracted in 96-well plates using the BioEcho EchoLUTION Plant DNA Kit (cat. #010-303-002). Libraries were prepared using the Twist 96-Plex Library Prep Kit (cat. #10653). Sequencing was conducted on an Illumina NovaSeq 6000 system at Discovery Life Sciences (Huntsville, AL). To increase depth and approximate the coverage of long-read sequences, each short-read sample was sequenced twice. Long-read and short-read data were not normalized to identical depths to minimize potential biases from randomization.

### Analysis

A combination of open-source, custom, and proprietary analysis pipelines was used. Khufu, a proprietary analytical tool designed for trait mapping, was adapted for long-read sequencing despite being originally developed for short-read data. KhufuPAN, an adaptation of Khufu that uses a graph based reference, requires a Graphical Fragment Assembly (GFA) file that incorporates the genomes forming the pangenome alongside a well-assembled reference genome, designated as the null reference. This null reference is used to assign coordinates for SNPs and structural variants. The GFA file is best created using Cactus (24). The GFA is subsequently converted to produce a parental VCF file. A series of filters are applied to remove low quality alleles, low-quality variants, monomorphic variants, or variants exhibiting polymorphism only with the null reference genome. The resulting Filtered-Variant set is stored in a single folder for reuse in multiple applications, including KhufuPAN bootstrapping.

KhufuVAR, a Khufu sub-tool, is used for calling and filtering variants. Raw FASTQ files, whether from short- or long-read sequencing, are processed through Khufu-core (https://www.hudsonalpha.org/khufudata/) and mapped to the GFA file using VG Giraffe (38). Variants failing to meet a minimum depth cutoff are masked, and those overlapping with the Filtered-Variant set are extracted. Given that variant segregation depends on the population structure, an additional set of filters can be applied, including Minor Allele Frequency (MAF), polymorphism, and missing data thresholds.

To optimize computational efficiency, KhufuVAR operates in two stages. In the first stage, samples are processed in batches. Each batch is aligned with the Filtered-Variant set, and population filter metrics are calculated. In the second stage, the batches are combined, and the population filters are applied across the entire dataset. The final output is exported in Khufu panmap format, enabling the side-by-side presentation of SNPs, short indels, and large indels.

The TifRunner assembly version 2 (TRv2) served as the reference genome for both Khufu and open-source analyses and was obtained from PeanutBase.org. Eighteen parental lines were sequenced using PacBio HiFi platforms, and their genomes were assembled with Hifiasm (v. 0.19.8) (39) before being scaffolded using an in-house tool, Pteranodon, with TRv2 as a reference. Comparative analyses of short- and long-read data were conducted using two approaches: a linear analysis, which relied on a single reference genome, and a pangraph analysis, constructed from the genomes of the 18 parental lines. Linear Khufu was used for SNP detection, while KhufuPan identified both SNPs and indels. Pangraphs used TRv2 as the anchor reference, incorporating genomes from the following lines: C431, CB7, CC477, CC812, Florida07, GA12Y, Georganic, GPNCWS17, IAC322, ICG1471, Lariat, Marc1, NC94022, TifNV, TR, and York. KhufuPan combined de novo calls from Linear Khufu that were not detected by KhufuPan alone.

Khufu analyses consisted of mapping and genotyping with population-specific filters. Filtering parameters included minimum and maximum depth thresholds set at 1 and 3000, respectively. Additionally, SNPs with a minor allele frequency of less than 0.1 or missing in 75% or more of the lines were removed, and lines missing 90% or more of SNP calls were excluded. Khufu automatically filtered out sequences aligning to multiple loci, and alignment confidence was assessed by comparing the number of sequences before and after filtering. Sequencing depth, defined as the average number of times a position in the genome was sequenced, was calculated by Khufu. Genome coverage, the percentage of the genome sequenced at least once, was determined using minimap2 (v. 2.26) and samtools coverage (v. 1.16.1) (40), (41). Chromosome-level coverage was averaged to estimate whole-genome coverage. Gene space coverage was calculated using bedtools intersect (v. 2.28.0) with genome annotations for TRv2 downloaded from LegumeInfo.org.

Large structural variations were identified using PacBio’s pbmm2 and pbsv (v. 2.9.0) pipelines, with TRv2 as the reference genome. The functional effects of these structural variants were assessed using snpEff (v. 5.2c) (42). Analyses included both merged and individual VCF files. Correlation analyses were performed in RStudio (v. 2023.06.1+524) using the stats (v. package. All correlations employed the Pearson method via the cor.test function.

Selection accuracy assessment was carried out by evaluating how many variants within the region of interest were found in common in the query and the parental reference genome containing that QTL of interest where those variants are unique to the donor parent. Scores were determined by dividing the number of variants in common over total variants in the region.

A cost per insight analysis was conducted by assigning scores to different data types. Missing data received a score of 0, SNPs (1 bp) were scored 1, indels (2–1000 bp) received a score of 2, and large structural variations (>1000 bp) were scored 3. These scores were done based on impact. SVs over 1,000 bp can disrupt an entire gene so these were scored the highest. These scores were summed on a per-sample basis, and costs per sample were calculated based on laboratory resources and equipment used (details provided in the Supplemental File).

