## Supplementary material for "Long-Read Low-Pass Sequencing for High-Resolution Trait Mapping": Supplmental Figures

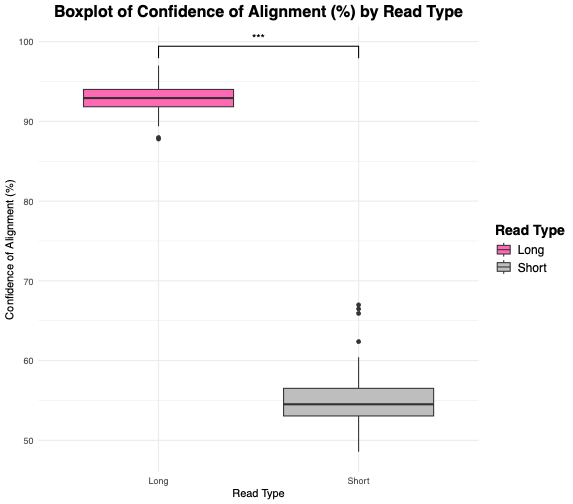

Supplementary Figure 1. A box plot comparing reads retained after filtering for misalignment, also called confidence of alignment. The long-read sequences show significantly higher confidence of alignment.

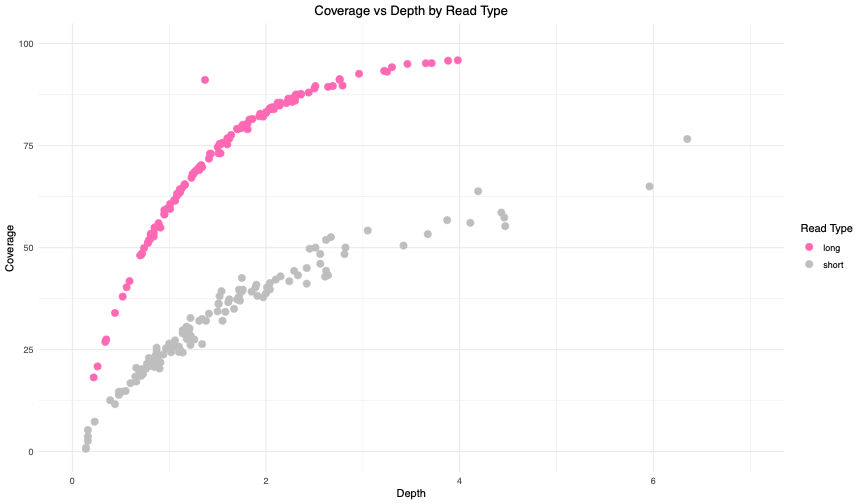

Supplementary Figure 2. A scatter plot showing the relationship between sequencing depth and genome coverage, with data separated by read type. Depth influences coverage to a point then plateaus (r=.56). Long reads are represented in pink, while short reads are shown in grey.

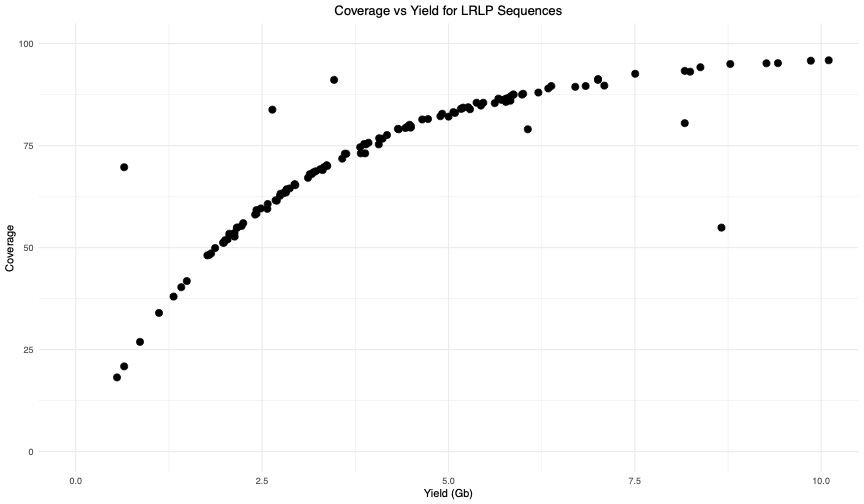

Supplementary Figure 3. A scatter plot showing the relationship between yield and coverage. A low yield is directly related to low coverage (r=.84).

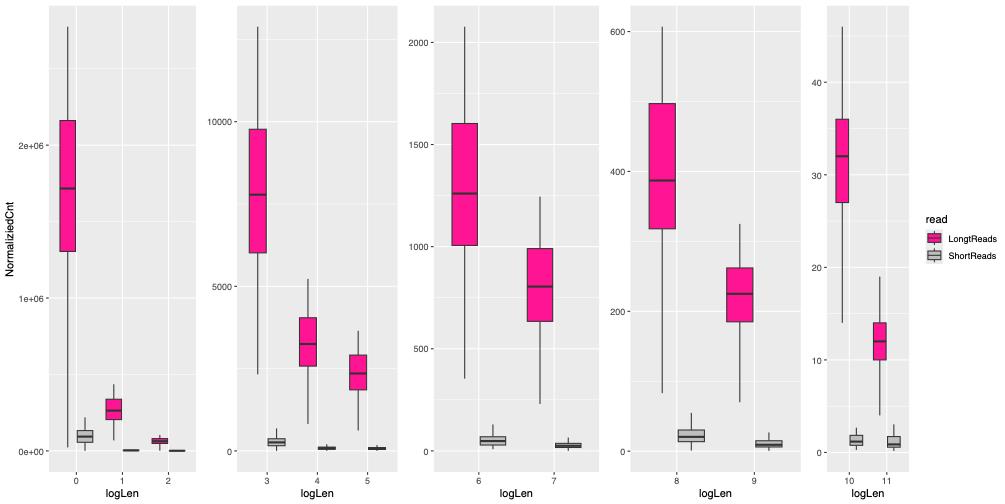

Supplementary Figure 5. A boxplot of structural variant (SV) lengths for long (pink) and short (grey) grouped by logarithmic values, where logLen group 0 represents SNPs and logLen group 11 represents SV’s above 50kb. In every group, long reads have much higher numbers of SVs.

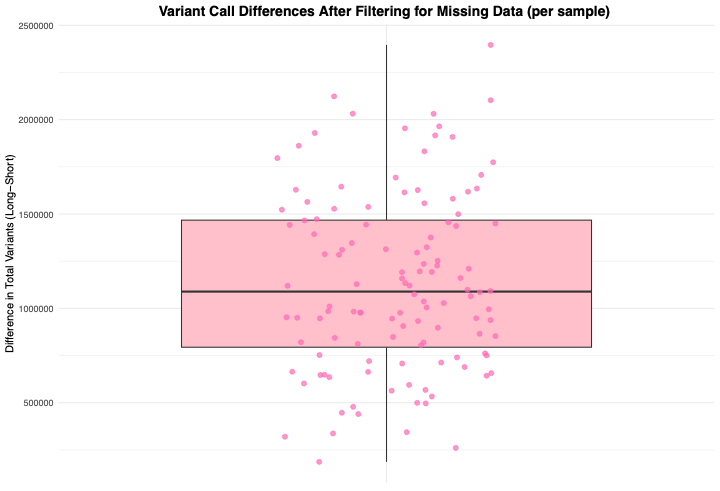

Supplemental Figure 4. A boxplot showing the difference in total variant calls between long read and short read data on a per-sample basis after filtering out missing data. On average, long reads called 1,137,853.725 more variants than short reads.

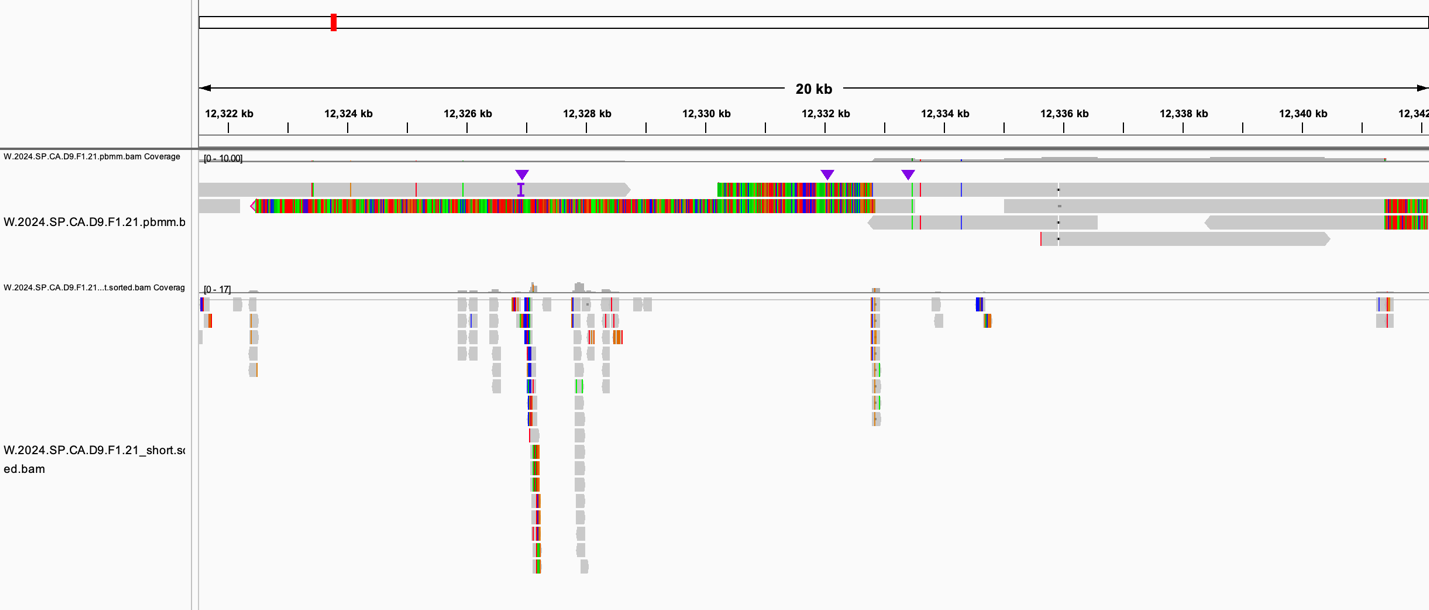

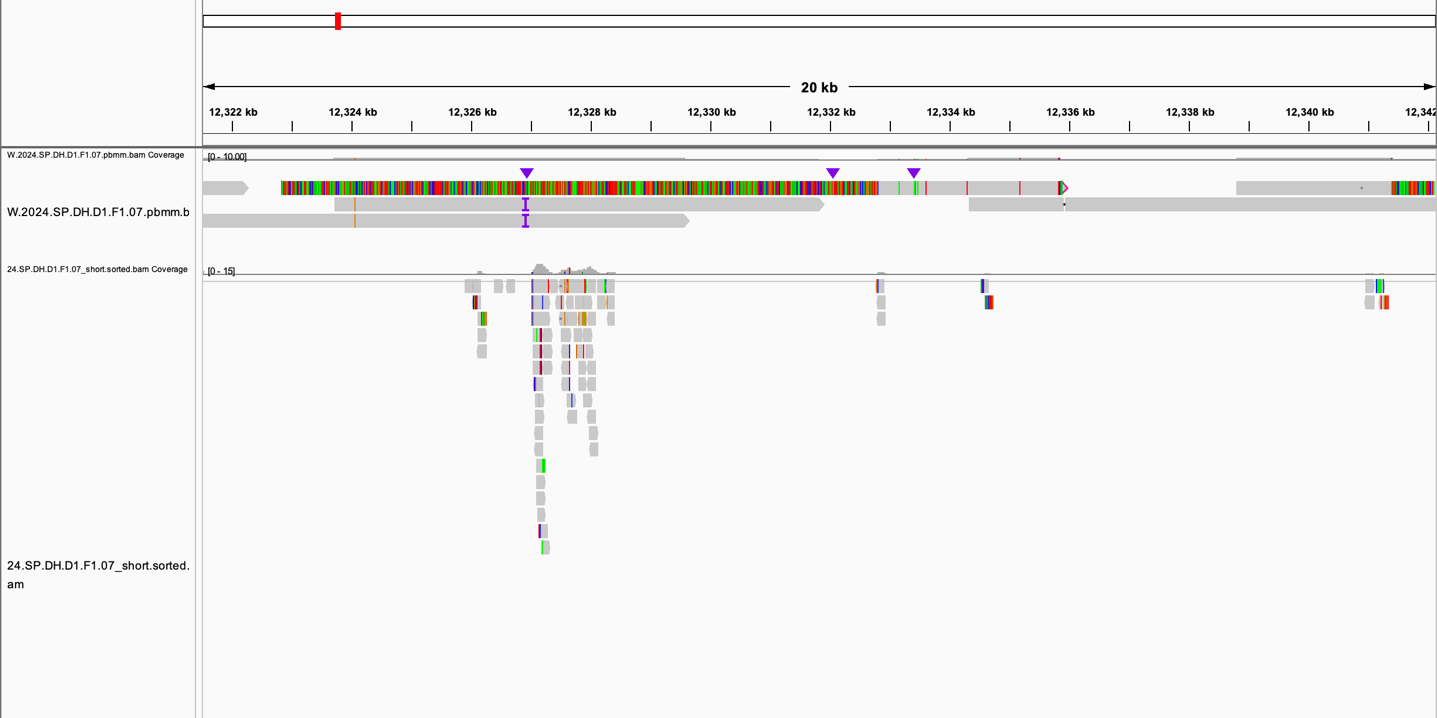

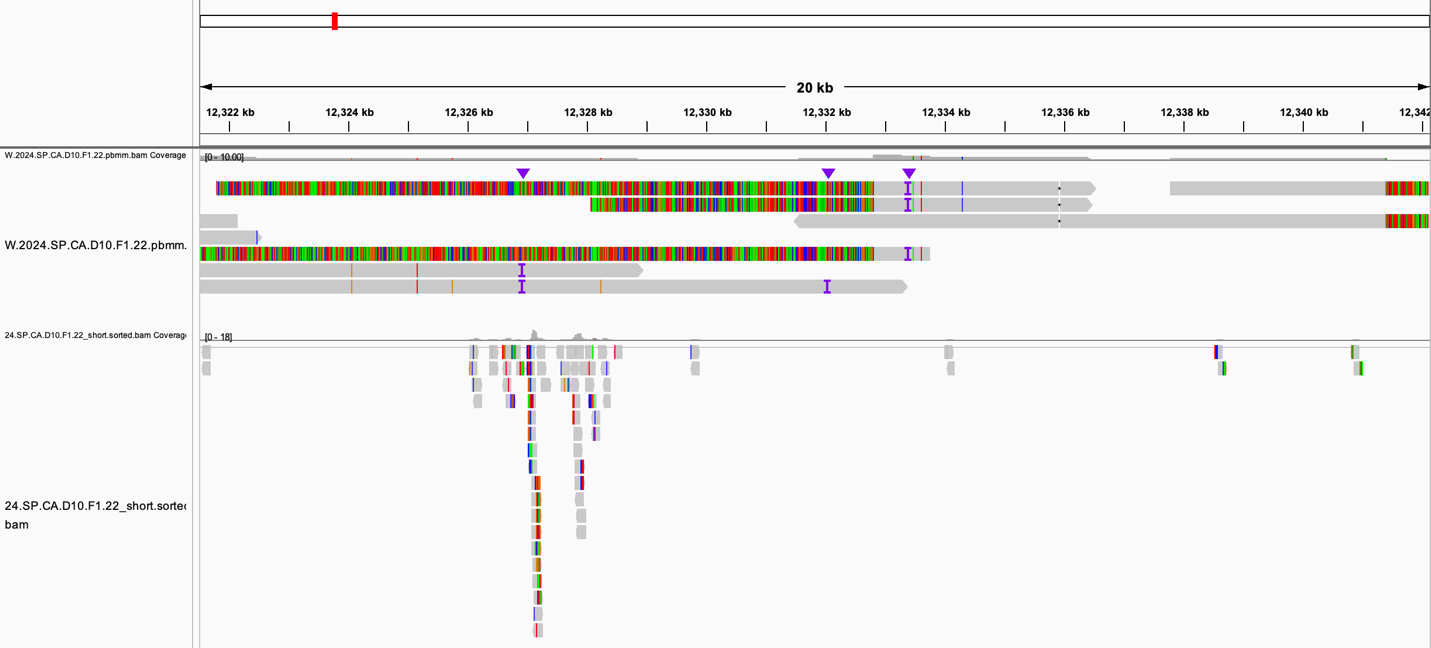

Supplemental Figure 5. Integrated Genomics Viewer (IGV) images of best scoring lines for TSWV similarity with LRLP data and the short-read counterparts. The IGV screenshot alignment of the respective reads to the region of TRv2 where the large insertion that confers resistance to TSWV maps. TRv2 contains one glutamate receptor gene in the region but lack the multiple copies that are found in the insertion, which confer resistance. The LRLP reads are on the top and show a long read with clipped reads spanning a 20kb region, covering the entire region. The partial clipping of the read indicates the insertion. Short reads are on the bottom and show some mapping to the region, but not enough to discover the TSWV disease resistance region insertion. Short reads are mapping to the edge of the insertion to one gene that is found in TRv2 but does not pick up on the additional genes which make up the insertion.
